# Zika and dengue viruses infecting wild-caught mosquitoes in an environmental protection area in Brazil

**DOI:** 10.1101/2019.12.17.879643

**Authors:** Karolina Morales Barrio-Nuevo, Antônio Ralph Medeiros-Sousa, Walter Ceretti-Junior, Aristides Fernandes, Iray Maria Rocco, Mariana Sequetin Cunha, Adriana Luchs, Luis Filipe Mucci, Renato Pereira de Souza, Mauro Toledo Marrelli

## Abstract

Species of the genus *Flavivirus* are widespread in Brazil and are a major public health concern. The city of São Paulo is in a highly urbanized area with some green spaces which are used for recreation and where potential vertebrate hosts and mosquito vectors of these arboviruses can be found, a scenario that can contribute to the transmission of flaviviruses to humans. This study therefore sought to investigate natural flavivirus infection in mosquitoes collected in the Capivari-Monos Environmental Protection Area (EPA) in the south of the city. Monthly mosquito collections were carried out from March 2016 to April 2017 with CO_2_-baited CDC light traps. Specimens were identified morphologically and grouped in pools. A total of 260 pools of non-engorged females were inoculated into the C6/36 cell lineage after analysis by indirect immunofluorescence assay (IFA). IFA-positive specimens were tested by qRT-PCR with genus-specific primers targeting a region of ~260 nucleotides in the flavivirus NS5 gene, and the PCR products were sequenced to confirm and identify the flavivirus species. *Anopheles* (*Kerteszia*) *cruzii* and *Wyeomyia* (*Prosopolepis*) *confusa* were the most frequent species collected. Zika virus (ZIKV) nucleotide sequences were detected in three mosquito species, *An. cruzii*, *Limatus durhami* and *Wy. confusa*, and dengue virus 2 (DENV-2) sequences in *Culex*. spp. and *Culex*. (*Mel*.) *vaxus*. To our knowledge, this is the first report of natural isolation of DENV-2 and ZIKV in sylvatic species of mosquitoes in the Capivari-Monos EPA. Our findings suggest that DENV-2 is present in *Culex* mosquitoes, and ZIKV in *Anopheles*, *Wyeomyia* and *Limatus*. The flavivirus species identified here are of medical importance; surveillance is therefore recommended in this EPA, where vertebrates and mosquitoes can act as flavivirus hosts and vectors.

## Introduction

Over 700,000 deaths worldwide every year are caused by infections transmitted by blood-feeding arthropods, accounting for 17% of all infectious diseases [1,2]. Mosquitoes have vector competence for viruses of great epidemiological importance, as seen in recent major outbreaks and epidemics of Chikungunya-virus (CHIKV), Dengue-virus (DENV), Zika-virus (ZIKV) and Yellow Fever-virus (YFV) infections in Brazil [3,4]. Arbovirus diseases occur worldwide, and their emergence and reemergence usually manifest as infections with mild to severe clinical symptoms in humans and domestic animals, occasionally progressing to death. These diseases therefore have a considerable impact on public health and the economy of the region affected [5–7].

Among the arthropod-borne viruses (arboviruses) circulating in Brazil, members of genus *Flavivirus* (family Flaviviridae) are noteworthy as they are the most common cause of viral infections and diseases in humans. In addition to DENV, ZIKV and YFV, several other flaviviruses of medical importance have been isolated in Brazil, including Bussuquara virus (BUSV), Cacicaporé virus (CPCV), Rocio virus (ROCV), Iguape virus (IGUV), Ilhéus virus (ILHV) and Saint Louis encephalitis virus (SLEV) [8–11]. Dengue is a reemerging disease in the country, with over 2 million confirmed cases and 702 recent deaths [12]. ZIKV has gained global attention as its geographic distribution has expanded dramatically from equatorial Africa and Asia to the Pacific Islands, South America and the Caribbean, causing many cases of neurological disorders and neonatal malformations [13–15].

Prevention and control of arboviruses require proper surveillance and vector control measures. Investment in appropriate integrated surveillance measures should therefore be a priority in Brazil, especially considering the size of the country’s population. Integrated surveillance, which covers epidemiological, entomological, sanitary and laboratory surveillance, is essential for the early detection of epidemics and for rapid, effective control measures [16]. Donalísio et al. [16] stress that investment in epidemiological, virological, vector and epizootic surveillance measures should be priorities in Brazil. In addition, in the absence of specific treatment and an effective vaccine, ongoing entomological and epidemiological surveillance should be strengthened and integrated to control and prevent these arbovirus diseases in Brazil [3].

The Capivari-Monos Environmental Protection Area (EPA), in the south of the city of São Paulo, Brazil, is a forest remnant close to urban areas. Previous studies in the Atlantic Forest in Brazil have shown that forest fragments offer favorable conditions for mosquitoes that are vectors of viruses to shelter and proliferate [17–19]. As circulating flaviviruses in city forest fragments can potentially cause disease outbreaks by infecting visitors and residents in neighboring areas, and given the lack of information about flavivirus-infected mosquitoes in parks in the city of São Paulo, the aim of this study was to investigate natural flavivirus infection in mosquitoes in the Capivari-Monos EPA and identify the virus species by nucleotide sequence analysis.

## Material and methods

### Study area and Mosquito Sampling

The study was conducted in the Capivari-Monos EPA, an area extending over 251 km^2^ in the Atlantic Forest in the extreme south of the city of São Paulo where sustainable use of natural resources is practiced (Fig 1). Representing around one-sixth of the area of the whole municipality and bordering on the Serra do Mar State Park, the area extends over the first hills and ocean slopes toward the upper reaches of Serra do Mar range, at altitudes varying from 740 to 800 m above sea level. It has a super-humid, tropical, ocean climate with average annual temperatures of around 19°C and rainfall of between 1,600 and 2,200 mm. The vegetation is dense, tropical, montane forest made up of Atlantic Forest remnants with different degrees of conservation, varying from well-conserved original forest to areas that have undergone a process of regeneration since 1950 and others that have been degraded recently as a result of rural and, especially, urban expansion. The district of Engenheiro Marsilac lies in the EPA and has around 10,000 inhabitants, most of whom are low-income settlers. The population density is approximately 41 inhabitants per km^2^ [20,21].

**Fig 1.**
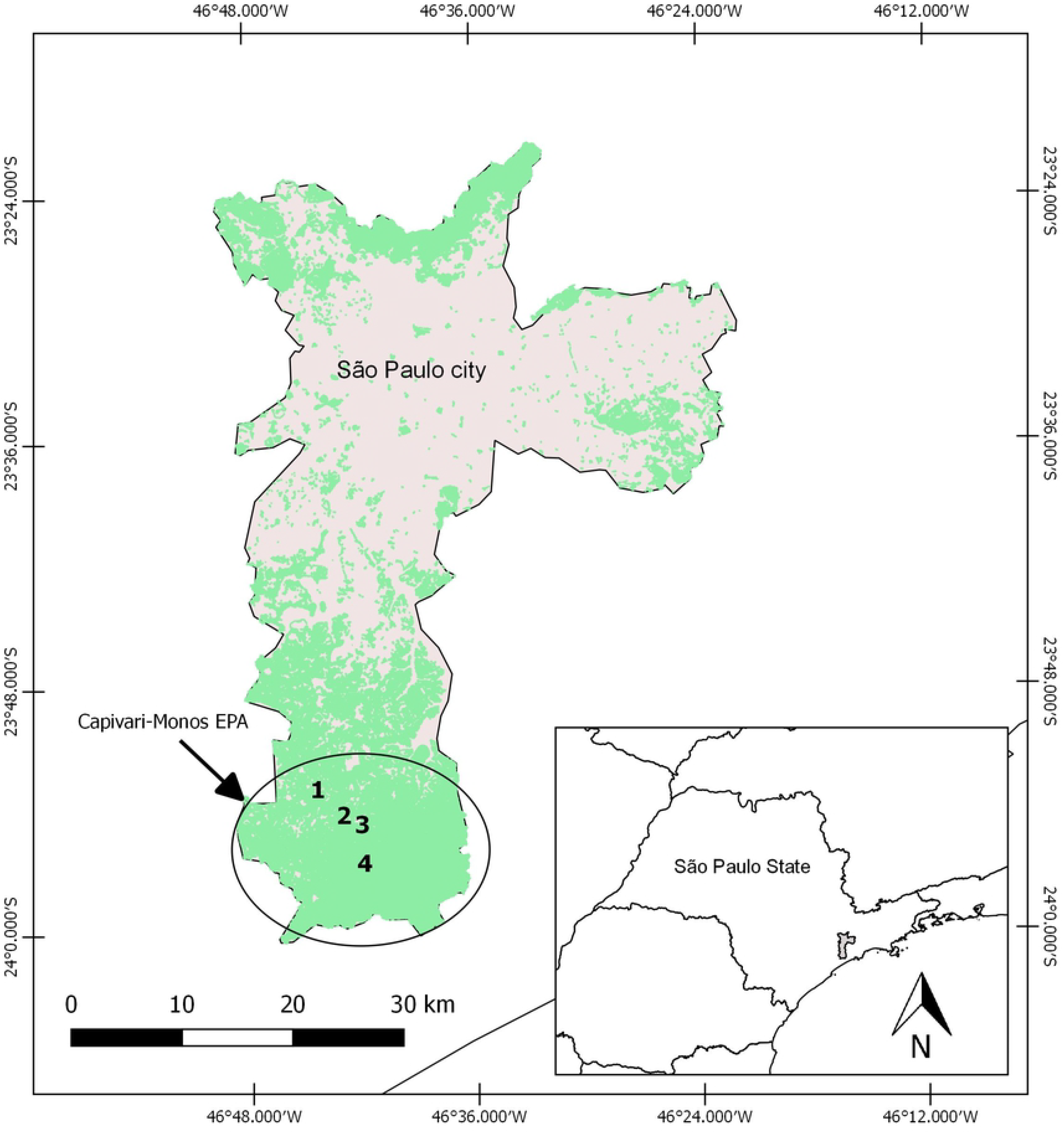
Capivari-Monos EPA in the south of the city of São Paulo, SP, Brazil. Collection sites are numbered as follows: (1) Embura village, (2) Marsilac village, (3) Transition zone, (4) Wild area. Green represents dense tropical forest; grey represents areas where there is human activity (villages, roads or rural properties).

Mosquito collections were carried out monthly from March 2016 to April 2017 in forested areas in Engenheiro Marsilac with different levels of anthropogenic intervention. Specimens were collected in the following areas: (1) Embura – a village surrounded by small farms and the EPA forest (23° 53.036′ S/46° 44.485′ W); (2) Marsilac – a village surrounded by the EPA forest and near a railway line (23° 54.395′ S/46° 42.486′ W); (3) Transition zone – private property near Marsilac village constituting a transitional area between a rural environment and the EPA forest (23° 54.556′ S/46° 42.167′ W); 4) Wild area – private property in the EPA forest next to a waterfall with a visitation area (23° 56.378′ S/46° 41.659′ W) (Fig 1). In each collection area two CO_2_-baited CDC light traps [22] with Lurex3™ were installed, one in the tree canopy (>10 meters) and another at ground level. All the traps were set up early in the afternoon and removed after 18 hours of exposure.

Specimens were carried alive to the Entomology in Public Health Laboratory at the School of Public Health, University of São Paulo (LESP/FSP/USP), where they were morphologically identified on a specially designed chill table with a stereo microscope and the dichotomous keys described by Consoli & Lourenço-de-Oliveira [23] and Forattini [24]. Non-engorged females were grouped in pools of up to 10 individuals according to their taxonomic category and place and date of collection. A total of 260 pools of mosquitoes were obtained in this way. The pools were then transported in dry ice to the Vector-borne Diseases Laboratory, Adolfo Lutz Institute, and stored at – 70°C until use.

### Detection of flaviviruses

Each pool was macerated in a tube containing 1 mL of 1.8 % bovine albumin and antibiotics (100 units/mL of penicillin and 100 µL/mL of streptomycin). The resulting suspension was centrifuged at 2500 rpm for 10 min, and the supernatant was collected and frozen at −70°C. Viral isolation was performed in cell tubes seeded with monolayer cultures of C6/36 cells (*Ae. albopictus* clone) containing 1 mL of modified Leibovitz’s (L-15) medium with 10 % fetal bovine serum (FBS), penicillin (100 units/mL) and streptomycin (100 µL/mL). Briefly, 100µL of the supernatant from each mosquito pool was inoculated into cell tubes after removing the medium. The cell tubes were then incubated for one hour at 28°C and slightly shaken every 15 minutes. A 1.5 mL volume of L-15 medium with 2 % FBS and antibiotics was then added, and the tubes were incubated for nine days at 28 °C. After incubation, the samples were analyzed by indirect immunofluorescence assay (IFA) [25] with the Saint Louis encephalitis anti-flavivirus polyclonal antibody produced at the Vector-borne Diseases Laboratory, Adolfo Lutz Institute.

### Identification of flaviviruses

Positive flavivirus samples were analyzed by quantitative reverse-transcription PCR (polymerase chain reaction) (qRT-PCR) and sequenced to identify the species. Viral RNA was isolated from 140 µL aliquots of the cell-culture supernatants with the QIAamp® Viral RNA Mini Kit (Qiagen, Valencia, CA, USA) according to the manufacturer’s instructions. The Pan-Flavi qRT-PCR assay previously described by Patel et al. [26] was run with the SuperScript® III Platinum® One-Step Quantitative RT-PCR System (Thermo Fisher Scientific, Waltham, MA, USA) according to the manufacturer’s instructions with genus-specific primers that target a 200-nucleotide region of the flavivirus NS5 gene.

The qRT-PCR amplicons (~260bp) were directly sequenced with the BigDyeTM kit v3.1 (Applied Biosystems, Inc., Foster City, CA, USA) and Flavi S and Flavi AS2 primers [26]. Dye-labeled products were sequenced with an ABI 3130 sequencer (Applied Biosystems, Inc., Foster city, CA, USA). Sequencing chromatograms were edited manually with Sequencher 4.7, and sequences were screened at the National Center for Biotechnology Information (NCBI) website with the Basic Local Alignment Search Tool (BLAST). The resulting sequences were edited manually and aligned with a set of cognate sequences of DENV and ZIKV available in GenBank (Supplementary Material Table S1) using Clustal W [27]. Minor manual adjustments to improve alignment were made with BioEdit 7.0.5.2 (Ibis Therapeutics, USA). Neighbor Joining (NJ) trees were constructed with the Kimura 2-parameter model determined by MEGA 6.0 with 1,000 bootstrap replicates [28]. Prototype ZIKV and DENV sequences (Table S1), available in GenBank, were added to the corresponding tree so that species identity could be confirmed.

Nucleotide sequences determined in this study were deposited in GenBank under accession numbers MK134005, MK134006, MK371391, MK371392 and MK371393.

## Results

In total, 878 specimens of Culicidae belonging to 37 taxa (11 genera) were sampled (Table S2), of which 99.8% were female and 0.2% male. Most of the specimens were collected in the canopy (54.1%), 41.1% were collected at ground level and for 4.8% information on stratification was not available. The species *An.* (*Ker*.) *cruzii*, (171 specimens), *Wy.* (*Prl*.) *confusa* (134), *Cx.* (*Cux*.) spp. (109), *Li. durhami* (61), *Cx.* (*Mel*.) *vaxus* (36) and *Wy*. (*Pho*.) *theobaldi* (58) were the most common species collected. More mosquitoes (61.4%) were collected in the wild area than in the transition zone (14.9%), Embura (13.1%) and Marsilac (8.7%).

qRT-PCR detected flavivirus RNA in 1.9% of the pools (5/260) of non-engorged females. RNAs from all the five flavivirus-positive pools were successfully sequenced, and the species were identified by comparing the 243-269 bp fragment of the partial NS5 gene with corresponding sequences from GenBank. DENV serotype 2 (DENV2) sequences were identified in *Cx.* spp. and *Cx. vaxus* pools, and ZIKV sequences in *An. cruzii*, *Li. durhami* and *Wy. confusa* pools. Flavivirus-positive samples were found in two areas in the EPA: wild area and transition zone (Table 1).

**Table 1.** Descriptions of flavivirus positive pools.

| Pool | Date | Location | Taxon | N | Stratification | Viral RNA | Fragment Size (bp) | Isolate | GenBank accession no. |
| --- | --- | --- | --- | --- | --- | --- | --- | --- | --- |
| 168 14P-F | Oct/16 | Wild area | *Cx. spp.* | 6 | Canopy | DENV-2 | 243 | Flavi-168-DENV-2 | MK134006 |
| 143 9P-C | Feb/17 | Wild area | *Cx. vaxus* | 4 | Canopy | DENV-2 | 255 | Flavi-143-DENV-2 | MK134005 |
| 148 11P-A | Feb/17 | Transition zone | *Li. durhami* | 6 | Ground | ZIKV | 242 | Flavi-148- ZIKV | MK371391 |
| 157 14P-A2 | Oct/16 | Wild area | *An. cruzii* | 10 | Canopy | ZIKV | 268 | Flavi-157- ZIKV | MK371393 |
| 150 12P-A | Feb/17 | Transition zone | *Wy. confusa* | 2 | Canopy | ZIKV | 269 | Flavi-150- ZIKV | MK371392 |

Descriptions of flavivirus positive pools according to name, collection date, collection site, taxon, number of specimens, stratification, viral RNA identified, fragment size, isolate and GenBank accession number.

The phylogenetic trees (Figs 2 and 3), which were constructed to confirm the species/serotype of the flaviviruses, include sequences from the isolates in this study (in bold) together with prototype sequences from GenBank obtained using BLAST.

**Fig 2.**
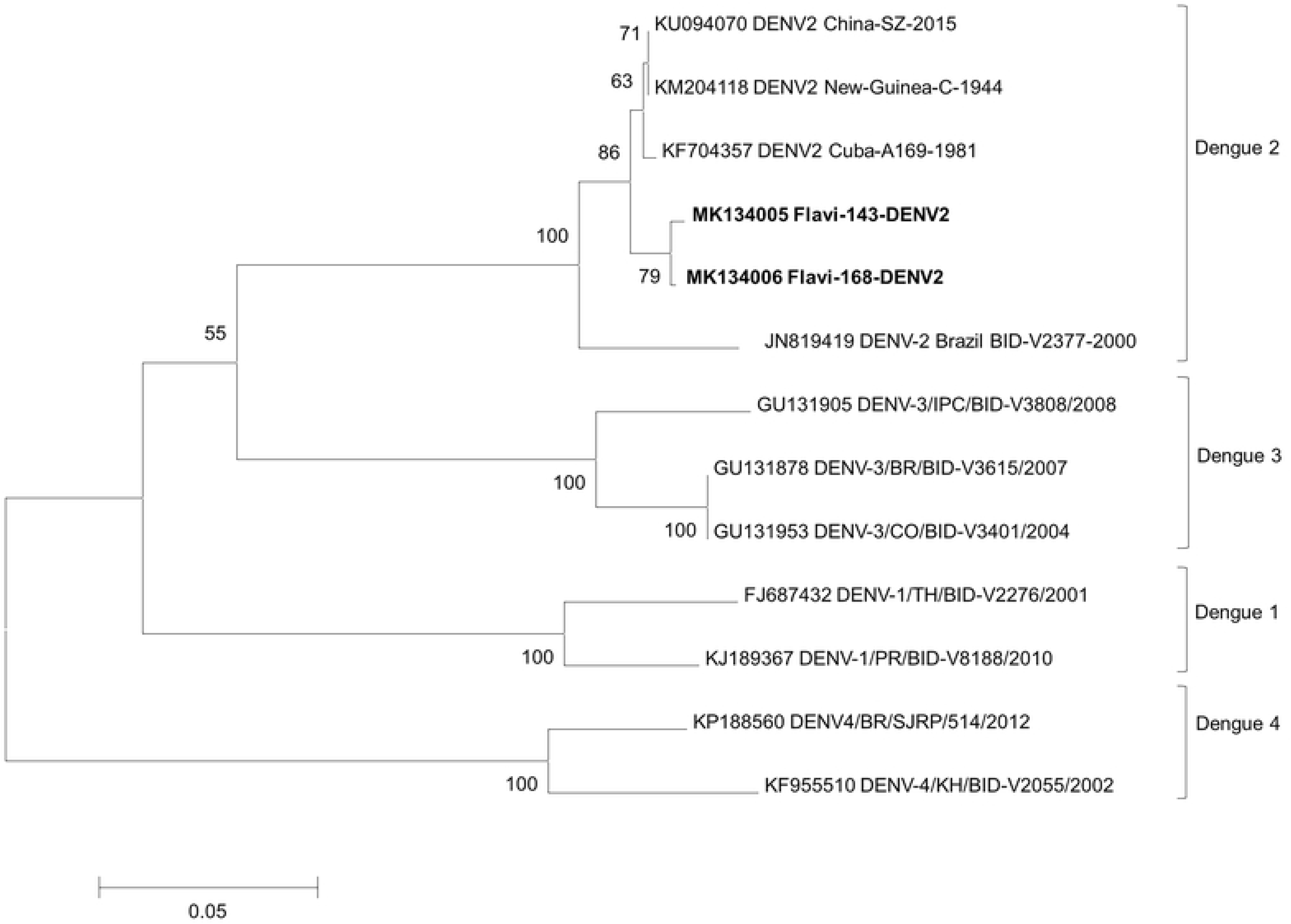
Neighbor-Joining (NJ) similarity tree. For the DENV-2 partial NS5 gene nucleotide sequences generated with MEGA 6.0. DENV-1, DENV-2, DENV-3 and DENV-4 sequences were obtained from GenBank. The species and accession number of each isolate are indicated. Table S1 gives information about each isolate. The scale indicates genetic diversity. The numbers at the nodes represent percentage bootstrap values.

**Fig 3.**
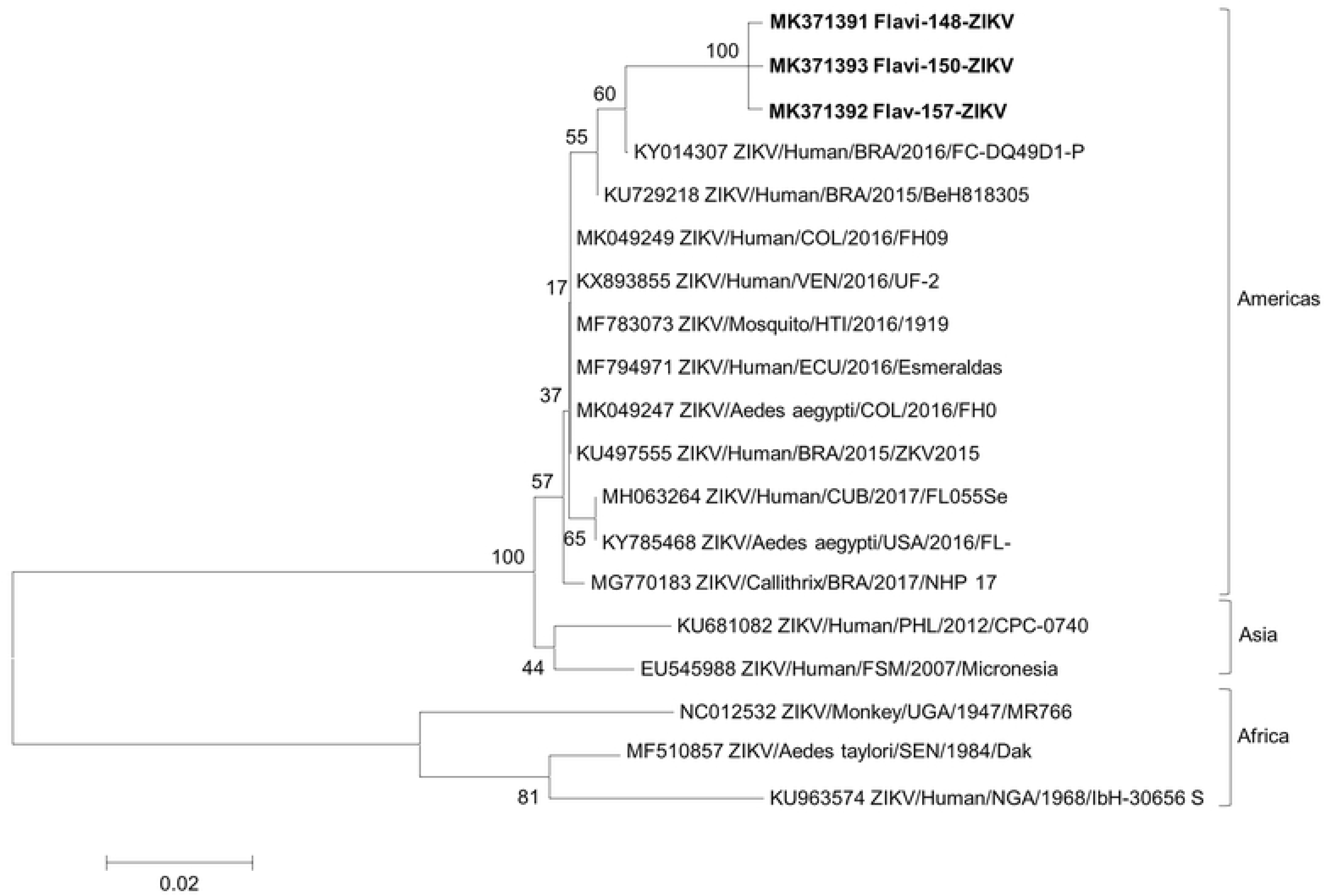
Neighbor-Joining (NJ) similarity tree. For the partial NS5 gene nucleotide sequences of the Flavi-148-ZIKV, Flavi-150-ZIKV and Flavi-157-ZIKV isolates generated with MEGA 6.0. Reference ZIKV sequences were obtained from GenBank. The species and accession number of each isolate are indicated. Table S1 gives information about each isolate. The scale indicates genetic diversity. The numbers at the nodes represent percentage bootstrap values.

The DENV tree confirmed the identification of isolates Flavi-143-DENV-2 and Flavi-168-DENV-2 as DENV-2 as they form a well-supported monophyletic group with the sequences from GenBank (Fig. 2). These two isolates showed a nucleotide (nt) identity of 94.1% with each other. Genetic analysis of the NS5 partial gene sequences revealed that the DENV-2 isolates identified in the present study had high nt similarity with human DENV-2 isolated in Cuba in 1981 (DENV2-Cuba-A169-1981), in China in 2015 (DENV2-China-SZ-2015) and in New Guinea in 1944 (DENV2-New-Guinea-C-1944) but low nt similarity with Brazilian human BID-V2377 isolated in 2000.

A nucleotide sequence identity of 99.6% was found when Flavi-148-ZIKV, Flavi-150-ZIKV and Flavi-157-ZIKV were compared with each other. Phylogenetic analysis of the partial NS5 gene sequences showed that these three isolates exhibited high nt similarity (97.0% to 98.1%) with other ZIKV viruses that have been isolated from humans, *Aedes* mosquitoes and non-human primates in the Americas since 2015. Three distinct ZIKV groups, from the Americas, Asia and Africa, can be observed in Fig. 3.

## Discussion

The finding reported here of sylvatic mosquitoes naturally infected with the DENV-2 and ZIKV flaviviruses for the first time in Brazil suggests that these two urban arboviruses are circulating in wild areas. Both DENV-2 and ZIKV originated in wild areas and settled in urban areas, causing a major impact on human health [29–31]. The presence of circulating flaviviruses in the Capivari-Monos EPA may indicate that an enzootic cycle involving vector mosquitoes and hosts (birds and mammals) has been maintained in this area and is likely contributing to transmission of these viruses, creating additional difficulties for disease control in the area.

The spread of arboviruses is directly related to the geographic expansion of their vector mosquitoes and vertebrate hosts and is very dependent on the virus-vector-host relationships [6,32]. Terzian et al. [33] detected ZIKV in non-human primates (NHPs) in urban and periurban areas in a city in the state of São Paulo and a few cities in the state of Minas Gerais, Brazil, suggesting that NHPs may be the vertebrate hosts responsible for circulation of ZIKV in these two states. The greatest diversity of Culicidae in Brazil is probably found in the Atlantic Forest [18], and several remnants of the Atlantic Forest can be found in various areas of the city of São Paulo, ranging from urban parks to conservation units. Conservation units farthest from the city center tend to be more preserved and consequently have greater mosquito diversity [18,34]. While the Capivari-Monos EPA is still a well-preserved area, it has recently undergone anthropic modifications that have led to fragmentation of the landscape [17,34].

Many studies have reported the presence of mosquitoes that are considered important flavivirus vectors in fragments of the Atlantic Forest in various areas in the city of São Paulo [18,34-36] and state of São Paulo [4,37,38], but natural flavivirus infection in these mosquitoes had not been detected prior to the present study. This is also the first report of mosquitoes naturally infected with flaviviruses in a conservation unit in the city of São Paulo. Unexpectedly, five species of mosquitoes (0.6%) not incriminated as flavivirus vectors were found to be naturally infected with species of genus *Flavivirus* in the study area. Most were collected in the canopy, including *An. cruzii*, *Cx.* spp., *Cx. vaxus* and *Wy. confusa*, but *Li. durhami* specimens were collected at ground level only.

*Culex vaxus* appears to retain behavioral characteristics typical of wild mosquitoes as it has been shown to adapt poorly to areas with reduced forest [39]. Information about this species is still quite scarce, and most of the available information is generally related to members of subgenus *Melanoconion*, to which this mosquito belongs. The species of this subgenus can develop in a wide range of breeding sites, from large natural water bodies, such as lakes, to small water retainers, such as bromeliads and artificial containers. They have eclectic feeding habits and can participate in natural arbovirus cycles [23–24]. Some species of the subgenus have been proven to be involved in the transmission cycle of arboviruses. *Culex* (*Mel.*) *pedroi* has been found naturally infected mainly by eastern equine encephalitis virus (EEEV) and western equine encephalitis virus (WEEV) and is a potential vector of Venezuelan equine encephalitis virus (VEEV) (*Alphavirus*). *Culex spissipes* is a potential vector for VEEV, Caraparu virus and other species of *Orthobunyavirus*. In addition, *Cx. portesi* may be involved in the Mucambo virus (*Alphavirus*) cycle in northeastern South America [24,40–42]. However, there have to date been no reports of *Cx. vaxus* infected naturally with species of genus *Flavivirus*. Of particular significance in the present study was the unexpected detection of *Cx. vaxus* naturally infected with DENV-2. Among dengue virus serotypes, DENV-2 is the most aggressive and most frequent circulating in Brazil and is currently responsible for 54.3 % of infections [43].

Mosquitoes of the *Culex* subgenus *Culex* develop in different breeding sites and feed on birds and mammals [24]. São Paulo metropolitan region is infested with *Cx. quinquefasciatus* as open-air breeding sites with high organic-matter content, such as the Pinheiros river, provide optimal conditions for the development and proliferation of this highly anthropophilic species, an opportunistic, cosmopolitan mosquito that has humans as its main host and feeds at night [44,45]. In the present study, DENV-2 was found in one of the positive pools of mosquitoes identified as *Cx.* spp. Although some species of genus *Culex* have been found to carry viruses in wild environments, Consoli & Lourenço-de-Oliveira [23] pointed out that the epidemiological importance of these species cannot be compared to that of *Cx. quinquefasciatus* in urban environments. To investigate ZIKV replication in *Cx. quinquefasciatus*, Guedes et al. [46] fed mosquitoes from Recife, Brazil, artificially and investigated the presence of ZIKV in their midgut, salivary glands and saliva. They demonstrated that *Cx. quinquefasciatus* is a competent vector for ZIKV, disagreeing with the findings of Lourenço-de-Oliveira et al. [47], who showed that these mosquitoes neither support replication of ZIKV nor play a role in transmission of this virus. Moreover, our study has demonstrated that other species of the *Culex* genus have vector competence for DENV-2. Hence, further studies of *Cx. quinquefasciatus* and other species of the genus are needed to clarify this issue.

*Anopheles cruzii* is generally restricted to the Brazilian coast, from the Northeast to South, following the original distribution of the Atlantic Forest. The high abundance of this species is directly related to the high availability of natural breeding sites. The species breeds mainly in epiphytic or terrestrial bromeliads, and the relative humidity on coastal slopes and plentiful shade under the forest canopy favor its adaptation to this environment. The abundance of this mosquito can also be attributed to its opportunistic behavior and eclectic feeding habits [23,24]. It is considered the species most associated with the transmission of human and simian *Plasmodium* in the Atlantic Forest, causing a type of malaria known as “bromeliad malaria”, so called because the immature forms of the mosquito develop in the water that accumulates in bromeliads. Although anophelines are usually known for transmitting malaria [50], some species have been found infected with arboviruses. Epelboin et al. [51] pointed out that *An. cruzii* is also responsible for transmitting O'nyong-nyong virus (*Alphavirus*), which is closely related to CHIKV. It is noteworthy that *An. cruzi* was also found naturally infected with the Iguape virus (a flavivirus) in 1994 in Juquitiba, SP [52]. Using artificial infection, Dodson & Rasgon [53] demonstrated that *An. gambiae* and *An. stephensi* are refractory to ZIKV infection. In addition, two species of genus *Anopheles*, *An. coustani* and *An. gambiae*, were found naturally infected with ZIKV in Africa [51]. However, there are to our knowledge no reports of *An. cruzii* naturally or artificially infected with ZIKV in the literature. This study can therefore be considered the first report of *An. cruzii* naturally infected with ZIKV in a conservation unit in the city of São Paulo.

The species *Wy. confusa* and *Li. durhami* belong to the Sabethini tribe and have close phylogenetic relationships. *Wyeomyia confusa* feeds during the day and is most commonly found at ground level but can also feed in the canopy. It is an opportunistic, eclectic species and frequently bites humans. Although it is usually sylvatic and breeds mainly in bromeliads, in this study *Wy. confusa* was observed in environments with greater anthropic interventions, such as Embura and the transition area, and was also found carrying ZIKV. Although this mosquito has already been found infected with arboviruses in wild environments, there is a dearth of information about its medical importance in the literature [24].

Even though *Li. durhami* is a sylvatic species, it is the species of the tribe Sabethini which best adapts to anthropic environments. It has been found frequently and abundantly in natural and artificial breeding sites that have undergone anthropic changes, usually where there is shady vegetation [23]. This mosquito bites actively throughout the day and feeds on birds and mammals at ground level. There have been reports of *Li. durhami* carrying the Guama, Tucunduba and Maguari viruses (*Orthobunyavirus*) [54]. Harbach [55] pointed out that although mosquitoes of the genus *Limatus* have been found with the *Wyeomyia* virus in Trinidad and the species *Li. flavisetosus* has been found with VEEV, it is unlikely that species of this genus are of epidemiological or even economic importance. However, in our study *Li. durhami* was detected carrying ZIKV. Although this is only the first such report worldwide, the importance of ZIKV for public health makes further studies of this species essential. It should be highlighted that the ZIKV identified here in three different mosquito species exhibited high nucleotide similarity with the other ZIKV isolates reported to date in the Americas.

Other mosquito species found infected with DENV include *Haemagogus leucocelaenus*(DENV-1), *Aedes albopictus* (DENV-3) [56], *Ae. aegypti* (DENV-1, DENV-2, DENV-3 and DENV-4) [57,58] and *Cx. quinquefasciatus* (DENV-4) [57]; species infected with ZIKV include *Ae. aegypti* [59, 60], *Cx. quinquefasciatus* and *Armigeres subalbatus* [60]. *Aedes aegypti*, considered the main vector of DENV and ZIKV in Brazil and other parts of the world [16,58], was not found in the Capivari-Monos EPA, probably because this species is an urban mosquito and the collection sites were in wild environments. *Aedes albopictus* was also sampled but tested negative for flaviviruses. The species is a potential vector of flaviviruses and many other arboviruses in some parts of the world [3,15]. Kucharz & Cebula-Byrska [61] note that *Ae. albopictus* can be a competent vector in regions where *Ae. aegypti* is not found in abundance.

Here we found sylvatic mosquitoes naturally infected with flaviviruses in an environmental protection area, the Capivari-Monos EPA. This finding may have implications for arbovirus surveillance programs as vector-borne DENV-2 and ZIKV transmission may occur in wild environments as well as urban areas. Further entomological surveillance studies are required to understand the potential role of these mosquitoes in maintaining the DENV-2 and ZIKV enzootic cycles and to elucidate how transmission cycles originated in the Capivari-Monos EPA would spread to other regions of the city. Surveillance activities typically include identification and laboratory confirmation of circulating arboviruses isolated from suspected cases of infection, as well as identification of the potential vector and monitoring of infestation rates for this vector. Reducing mortality due to flaviviruses depends on early detection of infection, an integrated surveillance system and a network for the provision of appropriate care and guidance during outbreaks and epidemics.

## Conclusion

This study reports the first isolation of flaviviruses from naturally infected mosquitoes collected in the Capivari-Monos EPA in the city of São Paulo, Brazil. *Culex.* spp. and *Cx. vaxus* were found infected with DENV-2, and *An. cruzii*, *Li. durhami* and *Wy. confusa* with ZIKV. These five mosquitoes were also the most abundant species collected. Considering the medical importance of *An. cruzii*, the main vector of bromeliad malaria in the Southeast of Brazil, it is imperative that further studies be carried out to investigate whether this species plays a role in ZIKV transmission. In addition, although there is to date no evidence that *Cx.* spp., *Cx. vaxus*, *Li. durhami* or *Wy. confusa* are of epidemiological importance, the fact that they were found infected with flaviviruses in this study means that they should be included in studies to investigate the flavivirus vector competence and capacity of these species and their potential to contribute to future dengue and Zika outbreaks and epidemics.

## Acknowledgments

We would like to express our gratitude to the following members of the field and laboratory teams at the Superintendency for the Control of Endemic Diseases, São Paulo Zoonosis Control Center, and the School of Public Health, São Paulo University: Dr. Ana Maria Ribeiro de Castro Duarte, João Carlos do Nascimento, Paulo Frugoli dos Santos, Luis Milton Bonafé, Antônio Waldomiro de Oliveira, Laércio Molinari, Gabriel Marcelino Neto, Luiz Sposito Jr, Renildo Souza Teixeira, Daniel Pagotto Vendrami, Laura Cristina Multini, Gabriela Cristina de Carvalho, Ramon Wilk da Silva, Rafael de Oliveira Christe, Eduardo Evangelista de Souza, Amanda Alves Camargo and Ana Leticia da Silva de Souza.

## Supporting information

**S1 Table. Reference sequences of dengue and Zika viruses from GenBank.** Sequences of dengue and Zika viruses from GenBank aligned to construct a phylogenetic tree by country of origin, isolate, year, origin of isolated material, genome sequence and GenBank accession number.

**S2 Table. Mosquito species collected in the Capivari-Monos Environmental Protection Area (EPA).** Mosquito species collected in the Capivari-Monos EPA according to the number of individuals at each study point, stratification, sex and number of pools. Mosquitoes collected from March 2016 to April 2017.

